# Viral Infection Detector: Ensemble Learning for Predicting Viral Infection in Single-cell Transcriptomics of Virus-Induced Cancers

**DOI:** 10.1101/2025.02.27.639934

**Authors:** Wenhao Han, Jiahui Hu, Kane Toh, Ming Ni, Roy Tan, Liang Wu, Xin Jin, Min Jian, Miao Xu, Hui Chen

## Abstract

Oncoviruses cause around 15-20% of all human cancers. As 3’ end-based single-cell RNA-sequencing (scRNA-seq) methods become more popular, the relationship between viral infection and tumor progression at the transcriptomic level has been increasingly studied at single-cell resolution. However, the identification of infected cells is challenging as viral reads from infected cells may not be captured, leading to high false negative rates. To recover these infected cells that have been missed, we developed VID, a stacked ensemble machine-learning model that predicts virus-infected cells in scRNA-seq datasets. Using VID, we uncovered biologically meaningful differences between infected and uninfected cells across different oncovirus-mediated cancers and different cell types. Our VID tool provides a user-friendly way to study changes induced by viral infections in scRNA-seq datasets.

## Introduction

Oncoviruses are estimated to cause approximately 15-20% of all cancers (Hausen and Villiers, 2014). According to the International Agency for Research on Cancer, 8 viruses are classified under the “carcinogenic” (group I) and “probably carcinogenic” (group 2A) categories: Epstein-Barr virus (EBV), Hepatitis B virus (HBV), Hepatitis C virus (HCV), Human Immunodeficiency virus type I (HIV-1), Human T-cell lymphotropic virus type I (HTLV-1), several high-risk Human papillomavirus (HPV) types, Kaposi sarcoma herpesvirus (KSH) and Merkel cell polyomavirus (MCV) (IARC monograph, 2012). Oncoviruses subvert a diverse set of host signalling pathways (e.g. PI3K-AKT-mTOR, MAPK, Notch, Wnt/ β-catenin and NF-κB), exploit the host DNA damage response system and evade the host immune surveillance, all of which contribute to their pro-tumorigenic capabilities (Ahmed and Jha, 2023; Krump and You, 2018).

Advances in single-cell technologies have empowered researchers to progress beyond the study of bulk, population-level averages towards the finer dissection of intratumor heterogeneity (ITH) and the role of the tumor microenvironment (TME) in viral carcinogenesis. (Cajal et al., 2020). In the context of nasopharyngeal carcinoma (NPC), Jin and colleagues performed full-length scRNA-seq (Smart-seq2) and 3’ droplet-based scRNA-seq (10x Genomics) on 19 Epstein-Barr virus (EBV) positive NPC and 7 non-cancerous nasopharyngeal samples. With these high-dimensional transcriptomic profiles at single-cell resolution, they uncovered a dual epithelial-immune meta-program associated with tumor cells with significant ITH. Malignant cells exhibiting a high score for this meta-program were associated with the co-expression of co-inhibitory receptors in CD8+ T cells, indicating their role in shaping the TME towards a pro-tumorigenic, immunosuppressive state. Studies on virus-induced cancers that utilise scRNA-seq (Wu et al., 2015, Cillo et al., 2020, Koya et al., 2021) have also highlighted the extensive heterogeneity that exist between cells or tissue samples, allowing researchers to distinguish between bystander and infected cells (Swaminath and Russell, 2024). By using two complementary full-length scRNA-seq technologies (mCEL-Seq2 and Smart-Seq2), Juhling and colleagues were able to quantify the HBV infection levels within single cells in hepatocellular carcinoma (HCC), which led them to discover a positive correlation between HBV-RNA levels and tumor cellular differentiation. Recently, spatial transcriptomic profiling of clinical samples has enabled researchers to chart the spatial distribution of transcriptionally active HBV integration events in liver biopsies of patients with chronic HBV infection (Yu et al., 2024), and to begin to investigate the relationship between EBV localisation and immune cell type composition in NPC (Liu et al., 2024). Thus, these technical breakthroughs, which lead to enhanced spatial and cellular resolution, lay the groundwork for the development of precision diagnostic tests and targeted therapies (Davis-Marcisak et al., 2021).

Despite recent improvements in single-cell technologies, several factors still make it challenging to accurately detect the expression of oncovirus transcripts within infected single cells. First, during the early stages of infection or the establishment of latency, viral transcript levels remain low within infected cells. Consequently, these low-abundance transcripts are likely to go undetected due to sampling variation that arise from the inefficiencies in RNA reverse transcription and biases in PCR amplification (Saliba et al., 2014). Additionally, human and virus-derived transcripts have differences in GC content, secondary structure and length of their poly(A) tails, which may further lower the RNA capture rate (Bost and Drayman, 2023). Whilst the problem of technical dropout can be addressed with specialised assays, such as a CRISPR-cas9 approach for the selective enrichment of HBV transcripts (Le et al., 2021), this approach necessitates additional costly experimental customisation. Third, existing popular single-cell RNA-seq platforms such as 10x Chromium tend to follow UMI tag-based rather than full-length based protocols, with most protocols capturing the 3’ ends of transcripts. Given the diversity and complexity of oncovirus transcript processing, reads derived from the 3’ end to a virus may not be distinct enough to differentiate between individual transcripts and isoforms (Depledge et al., 2019). For instance, all HBV virus transcripts share the same 3’ end, forming a nested set that comprise the HBx transcript sequence (Lamontagne et al., 2016; Seeger and Mason, 2015). The compact EBV genome expresses an extensive number of alternatively spliced genes, with dozens of isoforms identified across BHLF1, BZLF1 and the BART locus (Arvey et al., 2012). Similarly, the two pathogenic retroviruses, HTLV-1 and HIV, also rely heavily on alternative splicing in the synthesis of viral RNA products (Sertznig et al., 2018; Emery and Swanstrom, 2021). Fourth, whilst all oncoviruses apart from the Hepatitis C virus express polyadenylated RNA, several important transcripts are non-polyadenylated. The Epstein-Barr virus encoded small RNAs (EBERs) are small, non-coding, non-polyadenylated RNA which are abundantly expressed in the latent state, and thus serve as the gold standard molecular diagnostic tool for latent EBV infection (Abusalah et al., 2020). Recently, viral circular RNAs (circRNAs), which are non-polyadenylated, uncapped, covalently closed single-stranded RNA molecules, were identified in EBV, HPV and KSHV. circRNAs contribute to cancer pathogenesis by acting as miRNA sponges, regulating the rate of RNA transcription and protein synthesis (Tagawa et al., 2021, Sberna et al., 2023). As standard scRNA-seq protocols utilise poly-T-oligonucleotide primers for mRNA capture, non-polyadenylated transcripts are undetectable. Besides skewing the observed viral transcriptomic landscape, failing to detect highly expressed, non-polyadenylated transcripts may aggravate the false negative rates in single cell experiments.

Nevertheless, despite the dropout in viral reads, infected cells may display perturbations in their high-dimensional gene expression landscape relative to bystander cells. Machine learning algorithms excel in processing high-dimensional datasets with large numbers of cells, which would potentially allow them to identify relevant features to classify the infected cells from the bystander cells (Raimundo et al., 2021). In fact, simple machine learning classifiers may even outcompete more complex, deep-learning models. In a study by Abdelaal and colleagues, they compared the performances of 22 classification algorithms, ranging from classical methods like k-nearest neighbors and linear discriminant analysis to complex neural networks such as scVI (Lopez et al., 2018), and found that the support vector machine (SVM) with linear kernel had the best performance when tested across all 27 datasets (Abdelaal et al., 2019).

In this study, we leverage the predictive abilities of supervised machine learning tools via a stacked generalisation ensemble learning approach to classify the viral infection status of single-cells derived from 10x genomics 3’-end single-cell RNA seq datasets. Our tool provides a straightforward and easy-to-use computational solution to the task of viral infection status prediction without requiring any experimental modification to the standard library preparation workflow.

## Materials and Methods

### Description of scRNA-seq datasets

The Sorelle et al., (2022) cell line processed scRNA-seq dataset was downloaded from NCBI GEO under accession number GSE189141. the original publication. Raw scRNA-seq datasets for nasopharyngeal carcinoma (NPC) were downloaded from 4 NPC publications. Three of the raw datasets were downloaded from NCBI GEO under the accession numbers GSE162025, GSE150825, GSE150430, and another from the Genome Sequence Archive of the BIG Data Center at the Beijing Institute of Genomics under accession number HRA000087. Raw sequence data were then mapped to the human (GRCh38 GCF_002402265.1) and Epstein-Barr virus (ASM240226v1) genome references with CellRanger (version 5.1.0). The processed scRNA-seq dataset for the HPV oropharyngeal carcinoma (OPC) dataset was downloaded from NCBI GEO under accession number GSE182227.

### Downstream analysis of scRNA-seq datasets

Standard analysis for all datasets were conducted with Seurat v5.1.0 (Hao et al., 2024) in R. For the Sorelle et al., (2022) dataset, cells with number of read counts less than 25000, number of detected genes greater than 1000 and percentage of mitochondrial reads less than 20 were retained for further analysis. Pseudotime analysis was performed with Monocle3 (1.3.7). For the NPC datasets, cells with number of read counts greater than 1000, number of detected genes greater than 200, and percentage of mitochondrial reads less than 15% were retained for further analysis. Genes that were expressed in less than 3 cells were excluded from downstream analysis. Expression matrices were then normalized, and log transformed. Doublet removal was performed with DoubletFinder (v2.04). All samples were merged using the Seurat MergeData function. Harmony (v1.2.1)(Korsunsky et al., 2019) was used to remove potential batch effects. Top 2000 highly variable genes were selected for dimensional reduction. Principal Component Analysis was performed, retaining the top 25 principal components. The Leiden clustering algorithm was used for cell clustering and then projected into a 2-dimensional UMAP for visualization in reduced dimensions. ClusterTree (v0.5.1) was used to determine the optimal resolution for clustering. For the NPC datasets, to identify cell subtypes, each cell type was then performed a second round of clustering, batch effect removal and UMAP projection.as previously mentioned. We referred to the marker genes used in the original publications for cell type annotation. T cells subtypes were annotated according to Zhang et al. (2021), B cell subtypes were annotated following Ma et al., (2024), and monocytes subtypes were annotated following Chen et al., (2021). CellChat (v2.1.2) (Jin et al., 2025) was used for differential cell-cell communication analysis. For the Puram et al., (2023) OPC dataset, the processed data was analysed without incorporating additional filtering.

## Results

### Overview of Virus Infection Detector (VID)

VID is a supervised machine-learning classification pipeline that predicts the virus infection status of individual cells in labelled single-cell RNA-sequencing (scRNA-seq) datasets. Under the hood, it deploys a stacked generalization ensemble learning algorithm to learn meaningful gene expression patterns that distinguish between virally infected and uninfected cells.

#### Data requirements

To detect the virus infection status of single cells with VID, VID requires the scRNA-seq dataset to be labelled at the sample-level and cellular-level (Figure 1, Stage 1; Figure 2). The column in the metadata holding the sample-level labels should be specified using the “clinical_column” parameter, where samples are labelled as either ’positive’(infected) or ’negative’(normal). All cells derived from the normal samples are assumed to be uninfected and are classified as “negative”. These are treated as true negatives. Within the infected samples, VID further assesses the viral read counts at the single-cell level. If a viral read is detected within a cell, then the cell is classified as “positive” for infection and is considered a true positive. Otherwise, cells are classified as “unknown” and are the prediction targets for VID. Alternatively, users can manually provide the single-cell level labels in a text file and specify its directory with the optional parameter ‘label_dir’ during model training. With these labels, VID is trained on the true positive and true negative cells to learn gene expression patterns to identify the potentially infected cells (false negatives) within the set of “unknown” cells.

**Figure 1:**
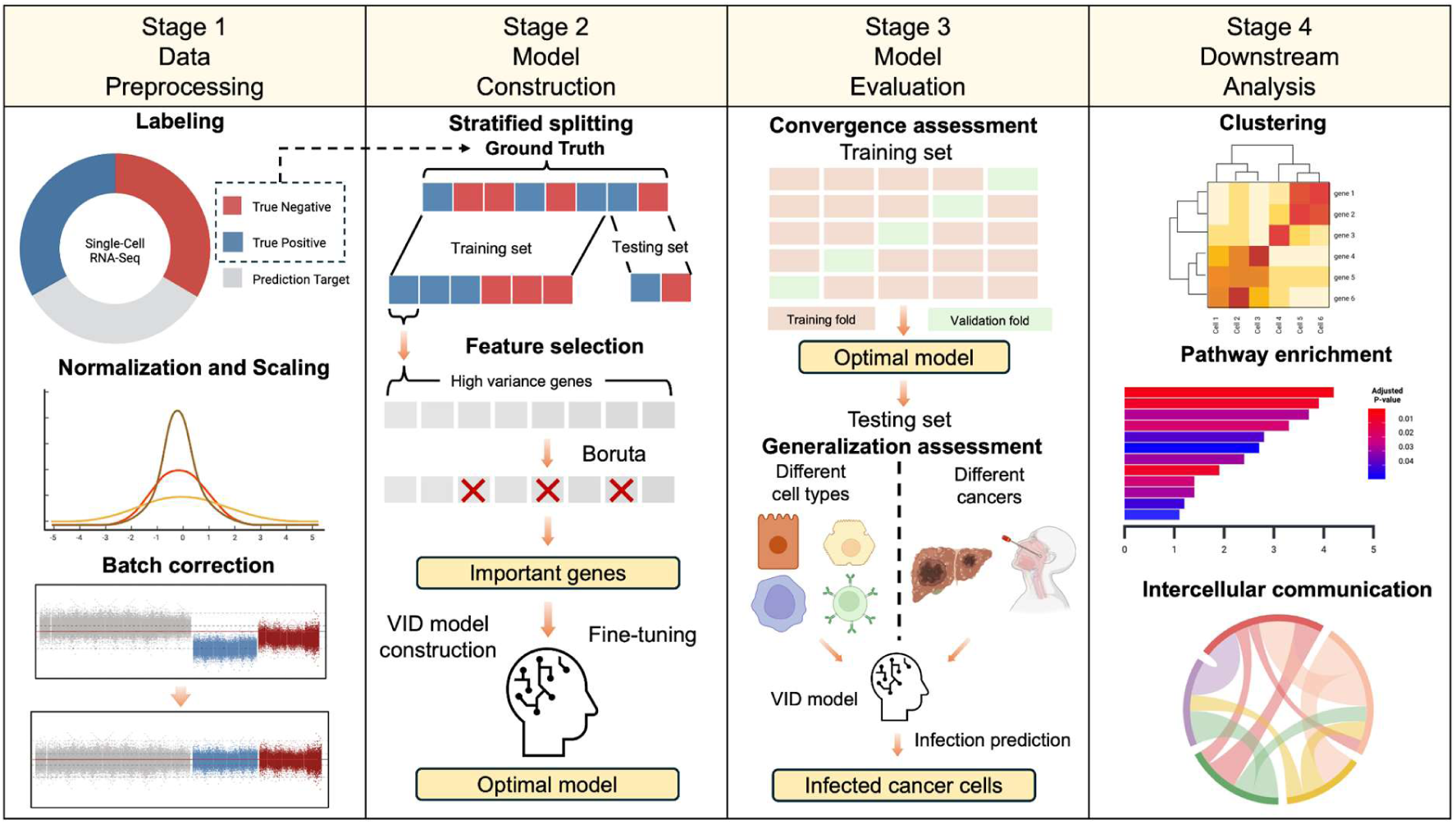
Overview of the VID workflow. The VID pipeline can be broken down into four stages.

**Figure 2:**
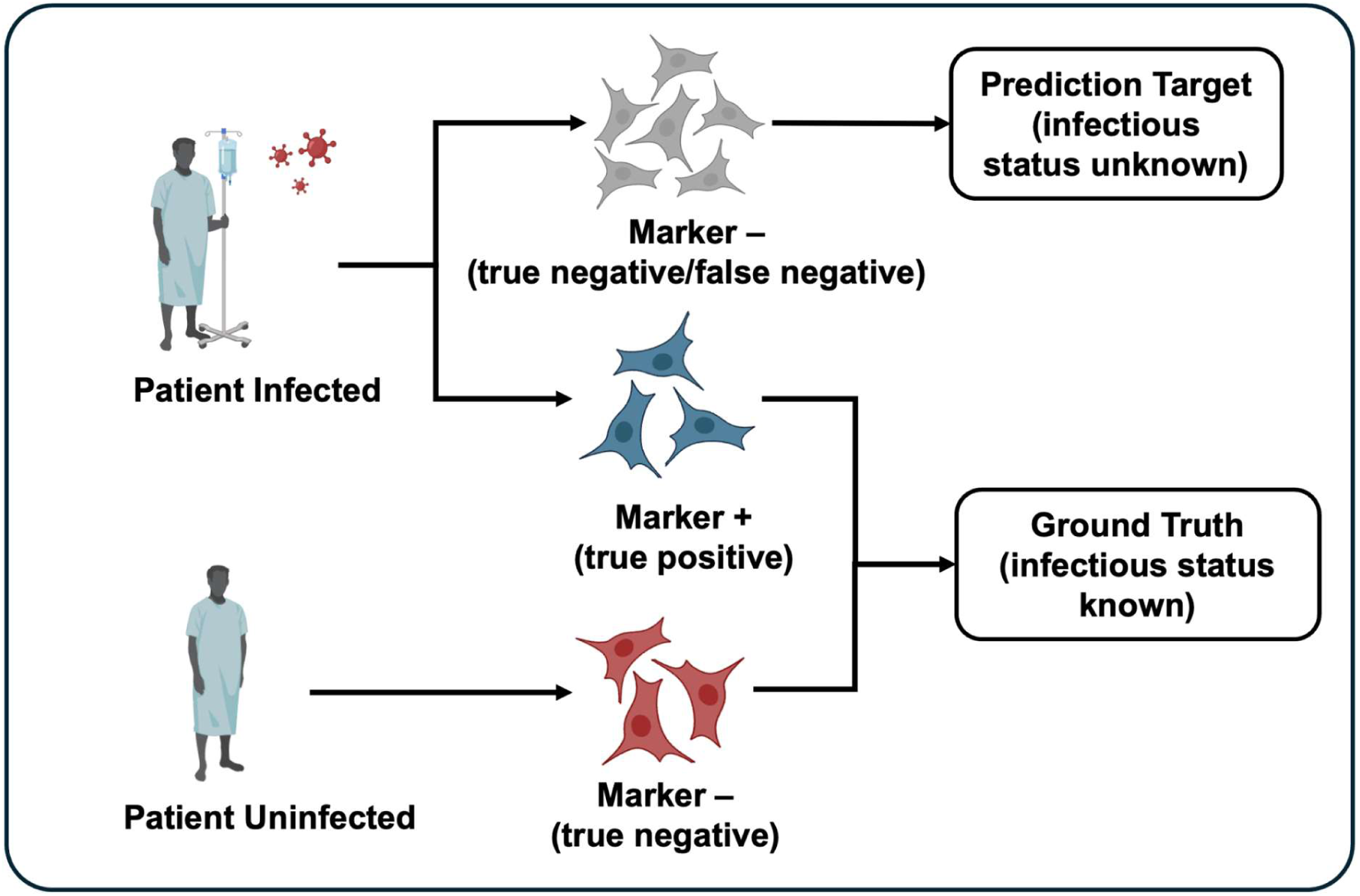
Data labeling at the sample and cellular level. Samples are labeled as either infected or normal (uninfected). Cells are labeled as either positive, negative or unknown, depending on the status of the sample that they are derived from and the level of detected viral reads. In particular, cells from infected samples that express a sub-threshold level of viral reads are classified into the “unknown” category, and are the prediction targets for VID.

#### Data preprocessing

The data preprocessing stages consist of normalization, scaling and batch effect correction. To stabilize the gene expression variance, log normalization is performed on the raw counts. To ensure the equal contribution of features and thus improve the performance of distance-based algorithms such as the k-nearest neighbors (KNN) and support vector machine (SVM) algorithms, VID applies z-score normalization to scale the log-normalized gene expression matrix. Crucially, systematic technical differences may arise from data collection across different timepoints or platforms, and these would confound model training and result in biased predictions. To address this, VID applies Harmony (Korsunsky et al., 2019) for batch correction in the final step of data preprocessing.

#### VID machine learning classification pipeline

VID is a supervised, stacked generalization-based (Wolpert, 1992) ensemble learning model for classification. It is a meta-learning algorithm that uses several base learners (level-0 models) and integrates the predictions from these level-0 models with a meta-model (level-1 model) to make a final prediction. At the expense of added complexity, this hierarchical structure allows the model to make more accurate predictions by combining the strengths of the diverse base models. Specifically, the VID model is constructed with two layers: in the first layer, the random forest (RF), SVM, KNN, gaussian naïve Bayes (GNB), and logistic regression (LGR) are applied as base models. These five level-0 models are selected for their diverse approaches to learning thus encouraging complementary predictions. In the second layer, two level-1 meta-models, extreme gradient boosting (XGB) and Multi-Layer Perceptron (MLP), are chosen. Users can choose the meta-model based on their requirements, with XGB being more interpretable and generally quicker to train, whereas MLP might offer a better performance in scenarios where the data exhibits deep hierarchical relationships and would thus benefit from a more complex learning structure.

We now describe the VID machine learning classification pipeline in greater detail (Figure 3). Following data preprocessing (See the Data Preprocessing section), to reduce the size of the dataset whilst preserving biologically meaningful information, VID first selects genes with the highest variance (HVGs) across cells. Intuitively, genes with low variance across cells are unlikely to differentiate between infected and uninfected cells. Next, VID performs stratified splitting to split cells into train and test sets. As opposed to random splitting, stratified splitting maintains a similar distribution of infected and uninfected cells across both train/test sets, avoiding class imbalance to encourage the model to generalize well to out-of-sample data. To avoid data leakage, in all analyses, viral markers are filtered out from the HVGs, retaining only the host genes in the dataset. Following this, VID further refines the set of informative features by applying the Boruta (Kursa et al., 2010) feature selection algorithm. The Boruta algorithm is a robust wrapper method that uses a RF model to iteratively assess the importance of each gene, identifying only those that are relevant for the prediction task. In this case, Boruta helps to focus the model on the most biologically significant genes for the viral status prediction task. This serves to reduce noise and overfitting, and by selecting a subset of HVGs, helps to further accelerate the subsequent training step.

**Figure 3:**
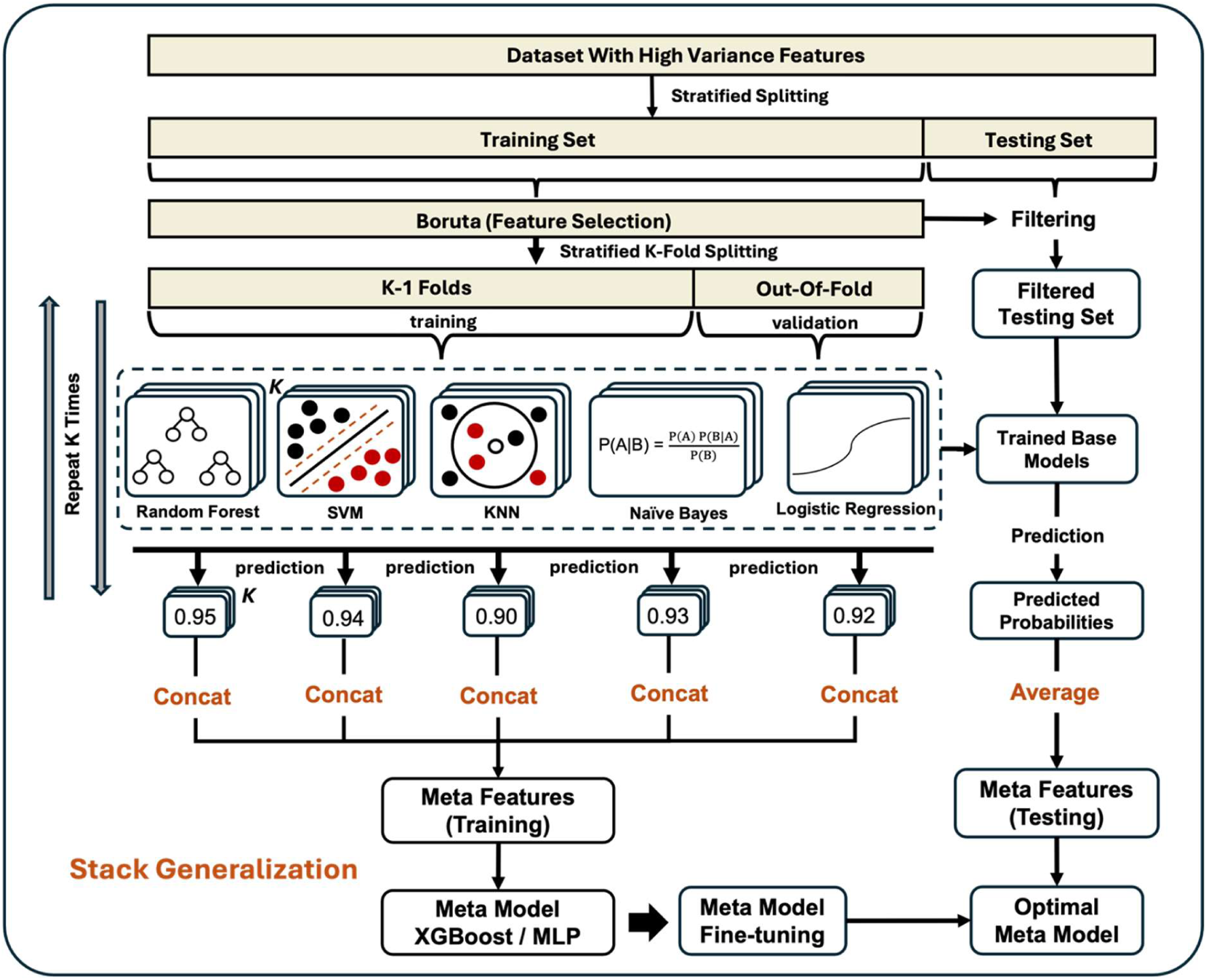
VID model architecture. The stratified K-fold cross validation stage is shown for the base model training stage, but omitted for the meta-model training stage for brevity.

After the Boruta feature selection step, stratified k-fold cross validation is carried out to split the training dataset into k equal-sized folds with similar proportions of infected and uninfected cells as the original dataset. For each of the K iterations, the 5 base learners are trained on K-1 folds, and validated on the remaining out-of-fold dataset. Let the out-of-fold dataset contain n cells. Then, K*n = *N* where *N* is the total number of cells in the training dataset. At the end of all K iterations, a total of 5*K*n out-of-fold prediction probabilities will be made, with each cell having 5 prediction probabilities, one from each base model. Users can choose between XGB and MLP for the meta-model. These out-of-fold prediction probabilities are concatenated to generate the meta-features which are fed into the chosen meta-model. The meta-model is fine-tuned as before with the same stratified splitting pattern, except that the meta-model input is now the 5 meta-features (instead of the gene expression features) for each cell.

The test dataset is filtered to contain only the genes chosen by the Boruta feature selection algorithm. Then, the 5 trained base models are applied to predict the probability of infection for each cell. To promote the generalisability of the meta-model, VID computes the mean of the K predicted probabilities for each base model as the input meta-feature for each cell and passes it into the trained meta-model. The trained meta-model uses these meta-features to make its final prediction regarding the probability of infection for each cell. For consistency, we use a threshold of 0.7 for all datasets in the publication - cells with a probability score equal or greater than 0.7 are defined to be predicted positive and are labelled as predicted negative otherwise.

#### Model performance evaluation

To ensure the generalisability of VID’s prediction capabilities, the test sets were derived from different disease samples and cell types (Figure 1, Stage 3). The accuracy, balanced accuracy, precision, recall, specificity, F1-score, and AUC metrics were computed to evaluate VID’s performance across the validation and test datasets. In particular, we computed a weighted average score for each metric to assess VID’s performance on imbalanced datasets. In addition, the confusion matrix and ROC curve were generated for the testing set to comprehensively summarize its performance and evaluate its robustness during classification. More advanced users may also find the provided CV and test scores to be useful for evaluating model convergence.

#### Downstream analysis

Following prediction (Figure 1, Stage 4), depending on their use case, users can set a threshold to classify cells as being either positive or negative for the virus infection. Users can then leverage this information to conduct further scRNA-seq downstream analysis. For example, they may visualize cell clustering results on the basis of this new grouping; run pathway enrichment analysis on the selected features to highlight host pathways that are perturbed upon viral infection, or investigate the changes in cell-cell communication patterns between uninfected and infected cells etc. We demonstrate the utility of these approaches in the results section of the manuscript. Users are encouraged to explore and implement more approaches that serve their specific analyses objectives.

#### VID software development

VID is written in Python and designed to be run through the command-line. It is open source and made freely available here: (https://github.com/HWHapply/VID.git).

### VID learns biologically meaningful features that distinguish EBV-infected and uninfected cells to improve the recovery of EBV-infected cells

In our first application, we assessed VID’s performance on the scRNA-seq dataset of primary B lymphocyte cell lines from Sorelle et al., (2022). Following quality control and the removal of T cells and CD14+ monocytes, the dataset contains 92,689 B cells grouped into 9 clusters (Figure 4A). Uninfected and post-EBV-infected cells show distinct changes in gene expression, with day 0 cells predominantly occupying clusters 2, 3, and 7, whereas infected cells are spread across the remaining 6 clusters (Figure 4B). To interrogate the importance of a viral prediction tool on this dataset, we first assessed the sparsity of the EBV viral read counts. Using a lenient thresholding strategy, we define a cell to be EBV-positive if it has at least 1 UMI count from the set of 91 EBV genes and label the cell as EBV-negative otherwise. 95% of the EBV-positive cells have a total of 15 EBV read counts or less across all EBV genes (Figure 4C). Most (38338 out of 39947) of all EBV-positive cells express the BCL2 homolog BHRF1, a lytic EBV gene expressed transiently during early infection to prevent immediate apoptosis (Altmann and Hammerschmidt, 2005). This is followed by EBNA-2, BWRF1, EBNA-1, and EBNA-3A, with the EBNA genes being expressed during latency. 33 of the EBV genes had no reads assigned in any of the EBV-positive cells. For the remaining 58 EBV genes that are expressed, around half (18845 out of 39947 EBV-positive cells) of all EBV-positive cells express BHRF1 exclusively, with significantly fewer cells expressing other viral genes in various combinations (Figure 4C). The skewed EBV expression pattern does not reflect the diverse EBV gene products typically expressed during infection (Damania et al., 2022). Furthermore, by examining the proportion of EBV-negative cells over the time-course study, we observe that even at the latest infection timepoint day 8, around 40% of cells remain EBV-negative (Figure 4D). Due to the high multiplicity of infection used in the study (Sorelle et al., 2022a) and the known sparsity in single-cell RNA-seq data (Bouland et al., 2023; Lähnemann et al., 2020), we therefore reasoned that the dataset likely contains a sizeable number of false negatives.

**Figure 4.**
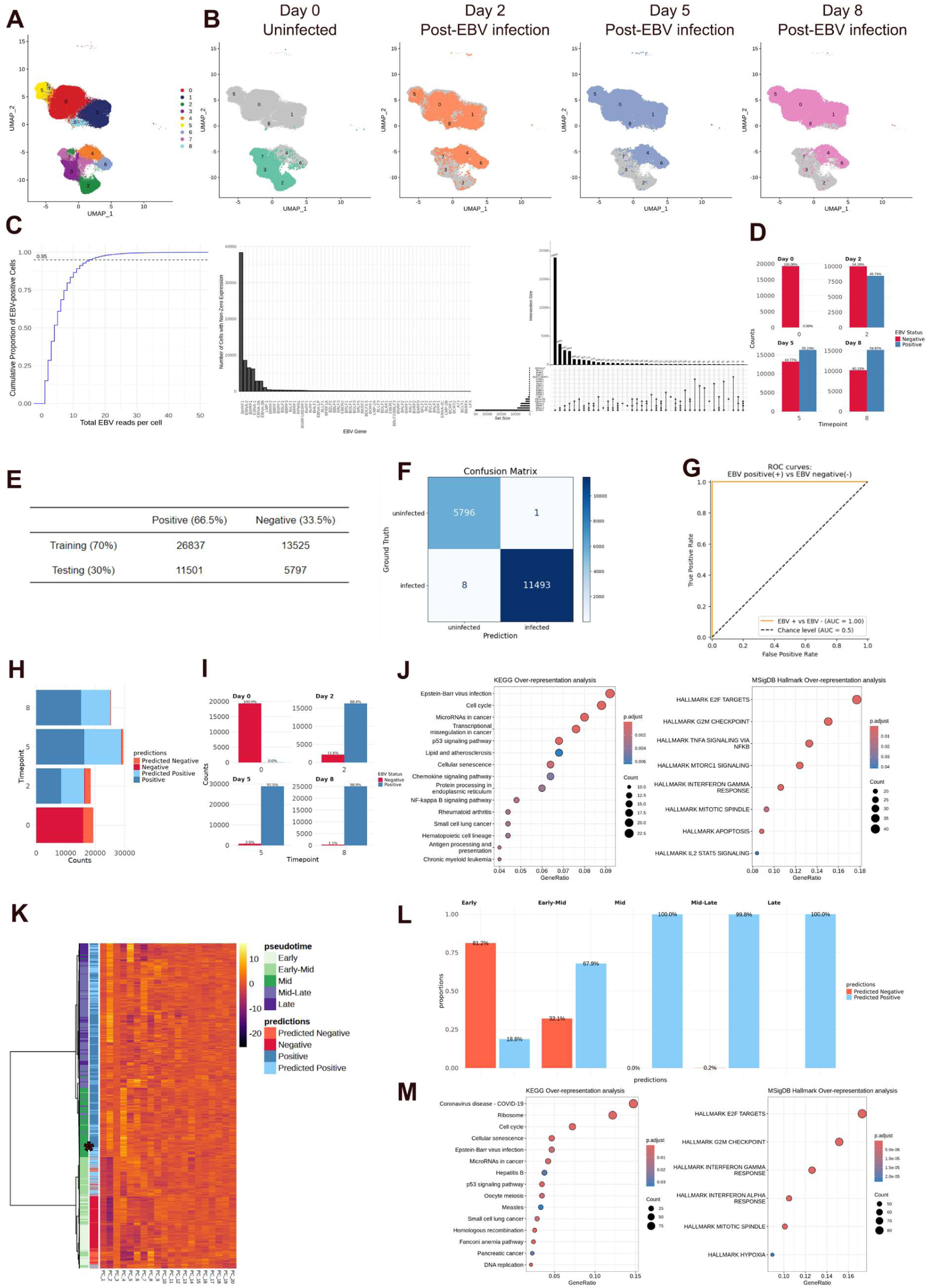
VID accurately recovers false negative EBV-infected cells in the time course single-cell RNA-seq dataset of pre and post EBV-infected primary B lymphocytes. A) Uniform manifold approximation and projection (UMAP) visualisation of the primary B cell single-cell RNA-seq dataset obtained from two donors, showing the labels for the 9 clusters. B) UMAP visualisation of the dataset, showing the distribution of cells from the uninfected (Day 0) and post-EBV infection timepoints (Day 2, Day 5, Day 8) across the 9 clusters. C) Evaluation of the sparsity levels of the EBV viral reads. EBV genes with non-zero expression levels (58 out of 91 EBV genes) were selected for display in the bar plot. The UpSet plot displays the intersection set patterns of the top 20 EBV genes with the highest read counts across all EBV-positive cells. D) Barplots showing the percentage of EBV-negative and positive cells across 4 timepoints before VID prediction. E) Distribution of EBV-positive and EBV-negative ground truth cell labels in the testing set. F) Confusion matrix for testing set predictions vs ground truth labels. G) ROC curve for VID’s performance. H) Horizontal barplots showing the distribution of prediction results on unknown cells and ground truth cell labels after VID prediction across the 4 timepoints. I) Barplots showing the percentage of EBV-negative and positive cells across 4 timepoints after VID prediction. J) KEGG and MSigDB Hallmark overrepresentation analysis of the features selected by the Boruta algorithm that distinguish between the EBV-positive and EBV-negative cells. K) Hierarchical clustering of the cells across the first 20 principal component embeddings. Cells are annotated according to VID predictions and their pseudotime scores. L) Percentage of EBV-positive and negative cells across the 5 pseudotime categories. M) KEGG and MSigDB Hallmark overrepresentation analysis of features enriched in early-mid predicted positive cells compared to early-mid predicted negative cells.

To test the classification performance of VID, we ran 70/30 stratified splitting of the dataset and evaluated the classification performance on the testing set (Figure 4E). In the testing set, only 1 out of 7597 of cells was a false positive and 8 out of 11501 cells were false negatives, and VID achieves a near-perfect AUC score (Figure 4F,G). After applying VID, almost all negative cells from timepoints 5 to 8 were predicted to be EBV-positive (Figure 4H). In comparison to the initial classification (Figure 4D), the percentage of EBV-positive cells at days 2, 5 and 8 have increased to 88.4%, 97.5% and 98.9% respectively, reflecting the increasing severity of viral infection across time (Figure 4I).

To verify that the predicted EBV-positive or EBV-negative cells were biologically meaningful, we assessed the quality of VID’s Boruta feature selection step. A total of 524 human genes were selected to classify the EBV-infection status. KEGG functional enrichment analysis indicate that these features are highly enriched in the Epstein-Barr virus infection pathway, as well as the cell cycle, microRNAs in cancer, p53 and NF-kB signalling pathways (Figure 4J). MSigDB Hallmark analysis revealed similar enrichment in the TNFA signalling via NFKB (Greenfeld et al., 2015) pathway, as well as the E2F and interferon gamma innate immune response pathways, consistent with a previous pan-cancer study of EBV infection (Song et al., 2019).

Next, we computed pseudotemporal analysis with Monocle3 using the uninfected cells from timepoint 0 as the anchor. We binned the pseudotime values into 5 discrete categories: Early, Early-Mid, Mid, Mid-Late and Late. Most of the predicted negative cells are derived from the Early and Early-Mid categories (Figure 4L) and generally, cells assigned to a later pseudotime value tend to have a positive or predicted-positive prediction (Figure 4K). Interestingly, 32.1% of cells in the Early-Mid category were predicted to be EBV-negative (Figure 4L), which is also reflected on the heatmap (Figure 4K). We sought to understand why VID gave these cells these predictions. To do so, we compared the predicted positive cells in the Early-Mid category against the predicted negative cells in the same category by performing functional enrichment analysis on the differentially expressed genes (DEGs). Functional enrichment analysis revealed pathways related to ribosome, EBV infection, E2F, interferon gamma and interferon alpha pathways (Figure 4M), highlighting that these predicted EBV-positive cells were in a more advanced stage of viral infection. Taken together, VID is able to identify informative features to recover the false-negative EBV-infected cells across various stages of the EBV-infection process.

### VID recovers infected epithelial and B cells in EBV-associated nasopharyngeal carcinoma cancer patient datasets

In our second application, we applied VID in real-world patient datasets for EBV-associated nasopharyngeal carcinoma (NPC) collected across 4 publications (Methods). Our combined dataset consists of a total of 325,810 cells across 22 cell types and 62 patients. (Figure 5A). As EBV displays dual tropism with the capacity to infect both epithelial cells and B cells (Borza and Hutt-Fletcher, 2002), we applied VID to predict the EBV infection status of both epithelial (13,416) cells and B (86,702) cells in the dataset. As before, we performed 70/30 stratified splitting (Figure 5C) and assessed VID’s performance on the test set across both cell types. VID achieves an AUC score of 0.97 and 0.98 for the epithelial and B cells respectively, correctly predicting the majority (260/271) of epithelial cells to be EBV positive and the majority (2162/2232) of B cells to be EBV negative (Figure 5D, E).

**Figure 5.**
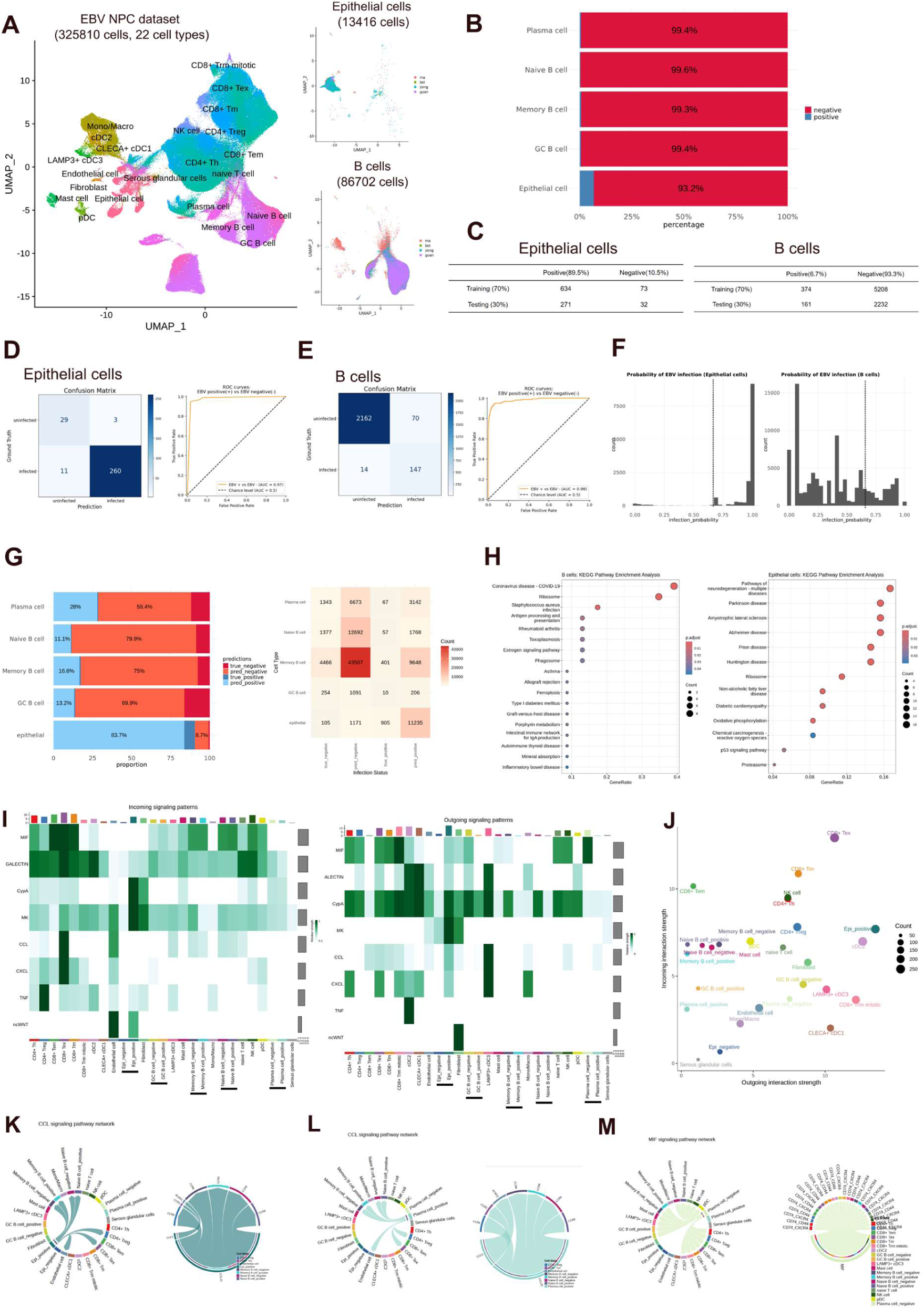
Applying VID to EBV-associated nasopharyngeal carcinoma datasets to predict EBV viral infection in epithelial and B cells. A) Uniform manifold approximation and projection (UMAP) visualisation of 325,810 cells from single-cell RNA-seq datasets collated across 4 publications. B) Barplots of the EBV positive and negative cells in the datasets prior to VID prediction. C) 70/30 stratified splitting of the epithelial and B cells in the dataset for testing. D-E) Confusion matrices and ROC curves for VID’s predictive performance for the epithelial and B cell test datasets. F) Probability of EBV infection for epithelial cells (left) and B cells (right). G) Percentage and number of predicted positive and negative cells in epithelial cells and B cell subtypes. H) KEGG overrepresentation analysis of the features selected by the Boruta algorithm that distinguish between the EBV-positive and EBV-negative cells in the B cells (left) and epithelial cell (right) populations. I) Heatmap with colors representing the relative incoming (left) and outgoing (right) signalling strength of signaling pathways across cell groups. The black bars highlight the epithelial and B cell subtypes. J) Scatter plot displaying the total outgoing or incoming communication probability associated with each cell group. Chord diagrams for the (K) CCL signalling pathway, showing signalling originating from both the Epi_positive and Epi_negative cell types and the ligand/receptors used during cell-cell communication. (L) CCL signalling pathway, showing signalling originating from both the EBV infected and uninfected plasma cell types and the ligand/receptors used during cell-cell communication. (M) MIF signalling pathway, showing signalling originating from both the EBV infected and uninfected plasma cell types and the ligand/receptors used during cell-cell communication.

Next, we leveraged VID to make predictions on the unknown epithelial and B cells (cells derived from EBV NPC cancer samples without any EBV viral reads present) in our dataset. For each unknown cell, VID outputs the probability that the cell is infected by the virus. The probability distribution differs for both cell types - the distribution for epithelial cells shows a strong left skew whereas the distribution for B cells appears flatter (Figure 5F). The genes identified by the Boruta algorithm for distinguishing EBV positive vs negative B cells are associated with the coronavirus disease and antigen processing and presentation pathways, whereas those identified in epithelial cells are associated with miscellaneous diseases and the dysregulation of metabolic pathways and oxidative phosphorylation (Figure 5H). This showcases VID’s capability to learn different probability distributions and different distinguishing features in a cell-type dependent manner.

Prior to VID prediction, most cells in the dataset are negative for EBV infection (Figure 5B). In stark contrast, following VID prediction, most epithelial cells in the dataset are predicted to be EBV positive (Figure 5G). Across the B cell subtypes, the predicted EBV-positive percentage is 11.1% for naïve B cells, 13.2% for germinal centre (GC) B cells, 16.6% for memory B cells and 28% for plasma cells. This is consistent with the dual tropism of the EBV virus in epithelial cells and B cells, where resting memory B cells act as a latent reservoir (Young and Dawson, 2014) which upon differentiation into plasma cells, lead to the initiation of viral replication and shedding (Calattini et al., 2010; Richardo et al., 2020).

Next, we examined the cell-cell communication patterns in the dataset, with a specific focus on comparing the differences in communication patterns between the EBV-infected and uninfected epithelial or B cell subtypes. We observed marked differences in the signalling patterns between the infected epithelial cell, plasma cell and GC B cell and their uninfected counterparts - the infected epithelial cell, uninfected plasma cell and uninfected GC B cell groups generally exhibit stronger incoming and outgoing signalling strengths (Figure 5I, J). Notably, Epi_positive cells show significant activity in the CXCL and CCL signalling pathways, whereas Epi_negative cells remain inactive in these pathways (Figure 5K, left). Specifically, the EBV-infected epithelial cells signal via the CCL5/CCR5 axis to the CD8+ exhausted T cells (Figure 5K, right). Whilst still under debate, our finding is consistent with recent reports of CCR5’s potential role in facilitating the recruitment of exhausted T cells and natural killer cells to shape an immunosuppressive TME (Aldinucci et al., 2020). CCR6 is predominantly expressed in B cells and CD4+ Tregs (Figure 5K, right), with evidence pointing to Tregs being induced by CCL20, leading to tumor invasion (Kadomoto et al., 2019). Interestingly, we find that the infected plasma cells are active in the CCL signalling pathway and signal via the CCL20/CCR6 axis to the same range of immune cell types as the EBV positive epithelial cells (Figure 5L). In contrast, the uninfected plasma cells are highly active in the macrophage inhibitory factor (MIF) signalling pathway and interact with a range of immune cell types via the CD74/CXCR4 axis (Figure 5M). In sum, by applying VID’s prediction to the EBV NPC datasets, we recovered a significant proportion of infected epithelial and B cells, which enabled us to gain greater insights into the biology of EBV NPC progression.

### VID achieves consistent performance in HPV-related oropharyngeal carcinoma

In our third application, we applied VID to an HPV-associated oropharyngeal carcinoma (HPV-OPC) dataset by Puram and colleagues (Puram et al., 2023). The dataset consists of 61565 cells across 14 annotated and 1 unresolved cell type (Figure 6A). HPV was detected only in the epithelial and unresolved cell types (Figure 6B, left), with most of the HPV-infected epithelial cells (6572) being derived from cancer samples (Figure 6C) of the atypical type (Figure 6D). Prior to VID prediction, around 54% of epithelial cells were HPV-negative (Figure 6E). After applying VID to the dataset, the percentage of HPV-positive cells (true positive and predicted positive cells) increased to around 70%, as an additional 24.6% of epithelial cells were predicted positive (Figure 6F).

**Figure 6.**
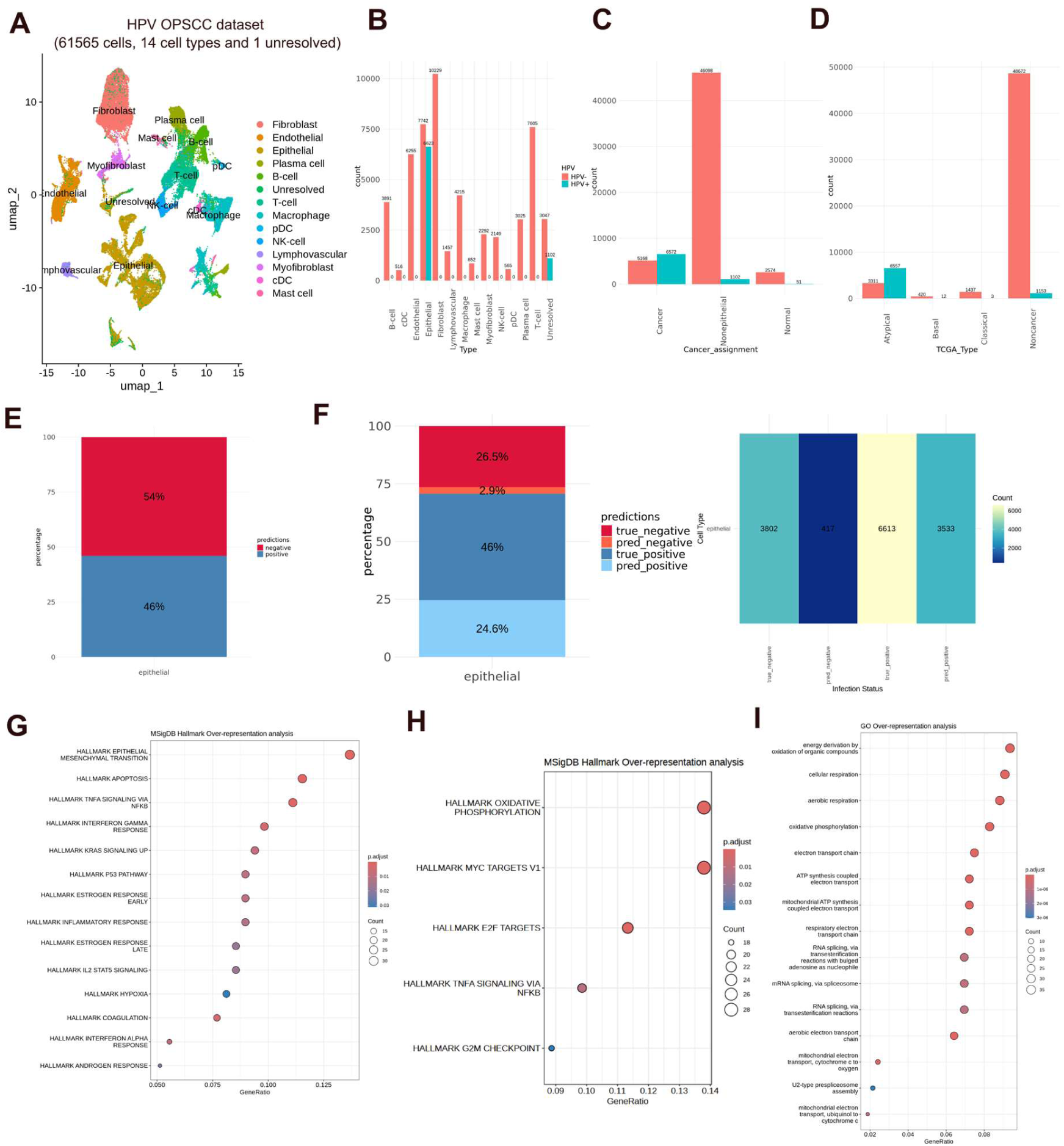
Applying VID to a HPV-related oropharyngeal cancer dataset to predict HPV viral infection in epithelial cells. A) Uniform manifold approximation and projection (UMAP) visualisation of 61565 cells from Puram et al., (2023). B) Barplots showing the distribution of HPV positive and negative cells across cell types. C) Barplots showing the number of HPV positive and negative cells across cancer assignment status. D) Barplots showing the number of HPV positive and negative cells classified by TCGA type. E) Percentage of HPV positive and negative epithelial cells prior to VID prediction. F) Percentage (left) and number (right) of HPV positive and negative epithelial cells after VID prediction. G) MSigDB Hallmark overrepresentation analysis of the features selected by the Boruta algorithm that distinguish between the HPV-positive and HPV-negative cells H) MSigDB Hallmark overrepresentation analysis of the genes that are upregulated in the predicted HPV infected cells relative to the uninfected cells. I) GO overrepresentation analysis of genes upregulated in the predicted HPV infected cells relative to the uninfected cells.

Over-representation analysis with the MSigDB hallmark gene set of the genes identified by the Boruta algorithm highlights association with known pathways relevant to HPV infection in OPC (Figure 6G). These include the process of epithelial mesenchymal transition (Saraf et al., 2023), proinflammatory TNFα signalling (Hong et al., 2020) and the modulation of apoptosis (Jamal et al., 2022). Next, to examine how VID makes its predictions, we examined the differentially expressed genes that are upregulated in the predicted HPV-positive epithelial cells relative to the predicted uninfected cells. Overrepresentation analysis of the MSigDB Hallmark and GO pathways (Figure 6H, I) reveals that the oxidative phosphorylation (OXPHOS) and aerobic respiration pathways are enriched, together with the E2F and G2M checkpoint processes involved in cellular proliferation. Our finding is consistent with recent evidence that the upregulation of mitochondrial oxidative metabolism activity through an increase in β_2_-Adrenergic receptor activity (HNSCC) is associated with a more aggressive phenotype (Lucido et al., 2018). In addition, metabolic dysregulation characterized by increased oxidative phosphorylation (OXPHOS) is associated with cellular proliferation, with NRF2 driving this proliferation in an OXPHOS-dependent manner in HPV-positive head and neck squamous cell carcinoma (Vyas et al., 2021). Taken together, we show that VID can be successfully applied to another different oncovirus-mediated cancer, facilitating the identification of key changes associated with viral infection and oncogenesis.

## Discussion

In oncovirus-mediated cancers studied with 3’ end-based scRNA-seq, detecting virally infected cells is challenging due to the low sensitivity of viral read detection, leading to a high rate of false negatives as virus-derived reads may not be expressed in infected cells. In this study, we introduce VID, a stacking ensemble machine learning model designed to improve the detection of infected cells in scRNA-seq datasets of oncovirus-induced cancers. To demonstrate the versatility and utility of VID, we evaluated it across three distinct scRNA-seq settings: (i) EBV-infected transformed lymphoblastoid cell line dataset (ii) integrated EBV-associated NPC datasets across 4 cancer patient studies (iii) an HPV-associated OPC dataset.

By using VID to recover the missing infected cells, we were able to detect biological differences distinguishing them from the uninfected cells. For instance, in the EBV-NPC datasets, 93.2% of epithelial cells were negative for EBV markers (Figure 5B). After applying VID, 83.7% were predicted to be EBV positive (Figure 5G). We were then able to identify activity in the CXCL and CCL signalling pathways that were concentrated in the EBV-positive infected cells. In the HPV-OPC dataset, we identified markers that were included in an 8-gene prognostic model (*Clorf105*, *CGA*, *CHRNA2*, *CRIP3*, *CTAG2*, *ENPP6*, *NEFH* and *RNF212)* of HPV-related squamous cell carcinoma of the head and neck (Mei et al., 2024). We identified 3 out of the 8 genes (CTAG2, NEFH and RNF212) as being more highly expressed in the predicted HPV-positive epithelial cells compared to the predicted HPV-negative epithelial cells. 1 of the 8 genes, ENPP6, was also identified but did not pass the significance test after Bonferroni correction for multiple testing. Therefore, VID can be generalised across oncovirus datasets and cell types to assist users in detecting biological differences between infected and uninfected cells.

Whilst VID provides a simple and valuable approach to identifying false negative virally infected cells, it has several limitations. Since our tool detects virus-infected cells based on their transcriptomic profiles, we may also classify some uninfected cells that display features associated with viral infection. This includes neighboring cells of virus-positive cells that have responded to cytokine inflammatory signals, as well as previously infected cells that retain a residual transcriptomic signature of viral infection. Consequently, it is more precise to describe our tool as identifying the cytokine activity and signalling pathways through which virus-positive cancer cells influence the TME. Nevertheless, the pronounced disparity between the frequencies of true-positive and predicted-positive cells observed in our analysis supports the notion viral infection rates are frequently underestimated in scRNA-seq data. Second, our default meta-model, XGB, does not consistently outperform all base models. For instance, on the EBV_EPI dataset, the LGR model achieves superior performance across all weighted evaluation scores. This outcome arises because the VID integrates three base models (RF, SVM, and LGR), where RF and SVM exhibit lower specificity (Supplementary Fig. 10 and Supplementary Table 2). This limitation is inherent to the meta-model, whose construction and optimization is guided by a specific metric (AUC). Alternatively, we can complement AUC with threshold-dependent metrics such as precision, recall, or the F1-score, to establish a more comprehensive evaluation standard for model construction. Another limitation of VID is its computational resource requirement, especially in the feature selection and model training steps, which becomes increasingly demanding for large datasets. We believe that in future iterations of VID, we can incorporate performance-based filtering or model pruning techniques to retain only the most effective base models, speeding up VID’s deployment. Assigning performance-based weights to base-models ensures that stronger models contribute more to the final predictions, balancing the impact of base models and maximizing the overall performance of the stacked ensemble. Fourth, VID requires mapped viral reads in at least a subset of cells to learn the gene expression patterns of infected cells. This requires the viral reference genome to be available and thus limits VID to being applied to known oncovirus-related cancers. We believe that VID can be used to augment the results of discovery-focused tools such as VirusPredictor (Liu et al., 2024) and viRNAtrap (Elbasir et al., 2023), thus extending its capabilities to previously undescribed viral infections.

In summary, we developed a novel ensemble-based method wrapped in a user-friendly software, VID, and demonstrated its utility across several scRNA-seq datasets. VID identifies virally infected cells in oncovirus-mediated cancers by combining the prediction outcomes of base learners and leveraging a meta-model (XGB or MLP) to make the final prediction. Our tool is especially useful for researchers who are interested to study the role of virus-infection in cancer progression, but whose analysis is unavoidably hindered by the sparsity of scRNA-seq transcriptomic datasets.

## Acknowledgements

We would like to thank DCS Cloud (https://cloud.stomics.tech/) for providing the computational resources and software support necessary for this study.

## References

Abdelaal, T., Michielsen, L., Cats, D., Hoogduin, D., Mei, H., Reinders, M. J. T., & Mahfouz, A. (2019). A comparison of automatic cell identification methods for single-cell RNA sequencing data. Genome biology, 20(1), 194. 10.1186/s13059-019-1795-z

Abusalah, M. A. H., Gan, S. H., Al-Hatamleh, M. A. I., Irekeola, A. A., Shueb, R. H., & Yean Yean, C. (2020). Recent Advances in Diagnostic Approaches for Epstein–Barr Virus. Pathogens, 9(3), 226. 10.3390/pathogens9030226

Ahmed K, Jha S. Oncoviruses: How do they hijack their host and current treatment regimes. Biochimica et Biophysica acta. Reviews on Cancer. 2023 Sep;1878(5):188960. DOI: 10.1016/j.bbcan.2023.188960. PMID: 37507056.

Aldinucci, D., Borghese, C., & Casagrande, N. (2020). The CCL5/CCR5 Axis in Cancer Progression. Cancers, 12(7), 1765. 10.3390/cancers12071765

Altmann, M., & Hammerschmidt, W. (2005). Epstein-Barr virus provides a new paradigm: a requirement for the immediate inhibition of apoptosis. PLoS biology, 3(12), e404. 10.1371/journal.pbio.0030404

Arvey, A., Tempera, I., Tsai, K., Chen, H. S., Tikhmyanova, N., Klichinsky, M., Leslie, C., & Lieberman, P. M. (2012). An atlas of the Epstein-Barr virus transcriptome and epigenome reveals host-virus regulatory interactions. Cell host & microbe, 12(2), 233–245. 10.1016/j.chom.2012.06.008

Borza, C., Hutt-Fletcher, L. Alternate replication in B cells and epithelial cells switches tropism of Epstein–Barr virus. Nat Med 8, 594–599 (2002). 10.1038/nm0602-594

Bost, P., & Drayman, N. (2024). Dissecting viral infections, one cell at a time, by single-cell technologies. Microbes and infection, 26(7), 105268. 10.1016/j.micinf.2023.105268

Bouland, G.A., Mahfouz, A. & Reinders, M.J.T. Consequences and opportunities arising due to sparser single-cell RNA-seq datasets. Genome Biol 24, 86 (2023). 10.1186/s13059-023-02933-w

Calattini, S., Sereti, I., Scheinberg, P., Kimura, H., Childs, R. W., & Cohen, J. I. (2010). Detection of EBV genomes in plasmablasts/plasma cells and non-B cells in the blood of most patients with EBV lymphoproliferative disorders by using Immuno-FISH. Blood, 116(22), 4546–4559. 10.1182/blood-2010-05-285452

Cheng, S., et al., A pan-cancer single-cell transcriptional atlas of tumor infiltrating myeloid cells. Cell,(2021). 184(3): p. 792–809 e23. doi: 10.1016/j.cell.2021.01.010.

Chen, Y. P., Yin, J. H., Li, W. F., Li, H. J., Chen, D. P., Zhang, C. J., Lv, J. W., Wang, Y. Q., Li, X. M., Li, J. Y., Zhang, P. P., Li, Y. Q., He, Q. M., Yang, X. J., Lei, Y., Tang, L. L., Zhou, G. Q., Mao, Y. P., Wei, C., Xiong, K. X., … Ma, J. (2020). Single-cell transcriptomics reveals regulators underlying immune cell diversity and immune subtypes associated with prognosis in nasopharyngeal carcinoma. Cell research, 30(11), 1024–1042. 10.1038/s41422-020-0374-x

Cillo, A. R., Kürten, C. H. L., Tabib, T., Qi, Z., Onkar, S., Wang, T., Liu, A., Duvvuri, U., Kim, S., Soose, R. J., Oesterreich, S., Chen, W., Lafyatis, R., Bruno, T. C., Ferris, R. L., & Vignali, D. A. A. (2020). Immune Landscape of Viral- and Carcinogen-Driven Head and Neck Cancer. Immunity, 52(1), 183–199.e9. 10.1016/j.immuni.2019.11.014

Damania, B., Kenney, S. C., & Raab-Traub, N. (2022). Epstein-Barr virus: Biology and clinical disease. Cell, 185(20), 3652–3670. 10.1016/j.cell.2022.08.026

Davis-Marcisak, E. F., Deshpande, A., Stein-O’Brien, G. L., Ho, W. J., Laheru, D., Jaffee, E. M., Fertig, E. J., & Kagohara, L. T. (2021). From bench to bedside: Single-cell analysis for cancer immunotherapy. Cancer cell, 39(8), 1062–1080. 10.1016/j.ccell.2021.07.004

Depledge, D.P., Srinivas, K.P., Sadaoka, T. et al. Direct RNA sequencing on nanopore arrays redefines the transcriptional complexity of a viral pathogen. Nat Commun 10, 754 (2019). 10.1038/s41467-019-08734-9

Elbasir, A., Ye, Y., Schäffer, D.E. et al. A deep learning approach reveals unexplored landscape of viral expression in cancer. Nat Commun 14, 785 (2023). 10.1038/s41467-023-36336-z

Emery, A., & Swanstrom, R. (2021). HIV-1: To Splice or Not to Splice, That Is the Question. Viruses, 13(2), 181. 10.3390/v13020181

Gong, L., Kwong, D.LW., Dai, W. et al. Comprehensive single-cell sequencing reveals the stromal dynamics and tumor-specific characteristics in the microenvironment of nasopharyngeal carcinoma. Nat Commun 12, 1540 (2021). 10.1038/s41467-021-21795-z

Greenfeld, H., Takasaki, K., Walsh, M. J., Ersing, I., Bernhardt, K., Ma, Y., Fu, B., Ashbaugh, C. W., Cabo, J., Mollo, S. B., Zhou, H., Li, S., & Gewurz, B. E. (2015). TRAF1 Coordinates Polyubiquitin Signaling to Enhance Epstein-Barr Virus LMP1-Mediated Growth and Survival Pathway Activation. PLoS pathogens, 11(5), e1004890. 10.1371/journal.ppat.1004890

Hong, H.S., Akhavan, J., Lee, S.H. et al. Proinflammatory cytokine TNFα promotes HPV-associated oral carcinogenesis by increasing cancer stemness. Int J Oral Sci 12, 3 (2020). 10.1038/s41368-019-0069-7

Jamal DF, Rozaimee QA, Osman NH, Mohd Sukor A, Elias MH, Shamaan NA, Das S, Abdul Hamid N. Human Papillomavirus 16 E2 as an Apoptosis-Inducing Protein for Cancer Treatment: A Systematic Review. Int J Mol Sci. 2022 Oct 19;23(20):12554. doi: 10.3390/ijms232012554. PMID: 36293403; PMCID: PMC9604055.

Jiaqiang Ma, et al., A blueprint for tumor-infiltrating B cells across human cancers. Science 384, eadj4857(2024). https://www.science.org/doi/10.1126/science.adj4857

Jin, S., Li, R., Chen, M. Y., Yu, C., Tang, L. Q., Liu, Y. M., Li, J. P., Liu, Y. N., Luo, Y. L., Zhao, Y., Zhang, Y., Xia, T. L., Liu, S. X., Liu, Q., Wang, G. N., You, R., Peng, J. Y., Li, J., Han, F., Wang, J., … Zeng, M. S. (2020). Single-cell transcriptomic analysis defines the interplay between tumor cells, viral infection, and the microenvironment in nasopharyngeal carcinoma. Cell research, 30(11), 950–965. 10.1038/s41422-020-00402-8

Jin, S., Plikus, M.V. & Nie, Q. CellChat for systematic analysis of cell–cell communication from single-cell transcriptomics. Nat Protoc 20, 180–219 (2025). 10.1038/s41596-024-01045-4

IARC Working Group Biological agents. IARC Monogr. Eval. Carcinog Risks Hum. 2012;100 Pt B:1–441.

Jin, S., Li, R., Chen, MY. et al. Single-cell transcriptomic analysis defines the interplay between tumor cells, viral infection, and the microenvironment in nasopharyngeal carcinoma. Cell Res 30, 950–965 (2020). 10.1038/s41422-020-00402-8

Jühling, F., Saviano, A., Ponsolles, C., Heydmann, L., Crouchet, E., Durand, S. C., El Saghire, H., Felli, E., Lindner, V., Pessaux, P., Pochet, N., Schuster, C., Verrier, E. R., & Baumert, T. F. (2021). Hepatitis B virus compartmentalization and single-cell differentiation in hepatocellular carcinoma. Life science alliance, 4(9), e202101036. 10.26508/lsa.202101036

Kadomoto, S., Izumi, K., Hiratsuka, K., Nakano, T., Naito, R., Makino, T., Iwamoto, H., Yaegashi, H., Shigehara, K., Kadono, Y., Nakata, H., Saito, Y., Nakagawa-Goto, K., & Mizokami, A. (2019). Tumor-Associated Macrophages Induce Migration of Renal Cell Carcinoma Cells via Activation of the CCL20-CCR6 Axis. Cancers, 12(1), 89. 10.3390/cancers12010089

Korsunsky, I., Millard, N., Fan, J., Slowikowski, K., Zhang, F., Wei, K., Baglaenko, Y., Brenner, M., Loh, P. R., & Raychaudhuri, S. (2019). Fast, sensitive and accurate integration of single-cell data with Harmony. Nature methods, 16(12), 1289–1296. 10.1038/s41592-019-0619-0

Koya, J., Saito, Y., Kameda, T., Kogure, Y., Yuasa, M., Nagasaki, J., McClure, M. B., Shingaki, S., Tabata, M., Tahira, Y., Akizuki, K., Kamiunten, A., Sekine, M., Shide, K., Kubuki, Y., Hidaka, T., Kitanaka, A., Nakano, N., Utsunomiya, A., Togashi, Y., … Kataoka, K. (2021). Single-Cell Analysis of the Multicellular Ecosystem in Viral Carcinogenesis by HTLV-1. Blood cancer discovery, 2(5), 450–467. 10.1158/2643-3230.BCD-21-0044

Krump, N. A., & You, J. (2018). Molecular mechanisms of viral oncogenesis in humans. Nature reviews. Microbiology, 16(11), 684–698. 10.1038/s41579-018-0064-6

Kursa, M. B., & Rudnicki, W. R. (2010). Feature Selection with the Boruta Package. Journal of Statistical Software, 36(11), 1–13. 10.18637/jss.v036.i11

Lähnemann, D., Köster, J., Szczurek, E. et al. Eleven grand challenges in single-cell data science. Genome Biol 21, 31 (2020). 10.1186/s13059-020-1926-6

Le, C., Liu, Y., López-Orozco, J., Joyce, M. A., Le, X. C., & Tyrrell, D. L. (2021). CRISPR Technique Incorporated with Single-Cell RNA Sequencing for Studying Hepatitis B Infection. Analytical chemistry, 93(31), 10756–10761. 10.1021/acs.analchem.1c02227

Liu, G., Chen, X., Luan, Y., & Li, D. (2024). VirusPredictor: XGBoost-based software to predict virus-related sequences in human data. *Bioinformatics (Oxford*, England), 40(4), btae192. 10.1093/bioinformatics/btae192

Liu, Y., He, S., Wang, XL. et al. Tumour heterogeneity and intercellular networks of nasopharyngeal carcinoma at single cell resolution. Nat Commun 12, 741 (2021). 10.1038/s41467-021-21043-4

Liu, Y., Ye, SY., He, S. et al. Single-cell and spatial transcriptome analyses reveal tertiary lymphoid structures linked to tumour progression and immunotherapy response in nasopharyngeal carcinoma. Nat Commun 15, 7713 (2024). 10.1038/s41467-024-52153-4

Lopez, R., Regier, J., Cole, M.B. et al. Deep generative modeling for single-cell transcriptomics. Nat Methods 15, 1053–1058 (2018). 10.1038/s41592-018-0229-2

Lucido, C.T., Callejas-Valera, J.L., Colbert, P.L. et al. β2-Adrenergic receptor modulates mitochondrial metabolism and disease progression in recurrent/metastatic HPV(+) HNSCC. Oncogenesis 7, 81 (2018). 10.1038/s41389-018-0090-2

Puram, S.V., Mints, M., Pal, A. et al. Cellular states are coupled to genomic and viral heterogeneity in HPV-related oropharyngeal carcinoma. Nat Genet 55, 640–650 (2023). 10.1038/s41588-023-01357-3

Raimundo F., Papaxanthos L., Vallot C., Vert J.-P. Machine learning for single cell genomics data analysis. Curr. Opin. Syst. Biol. 2021; 26:64–71.

Ramón Y Cajal, S., Sesé, M., Capdevila, C., Aasen, T., De Mattos-Arruda, L., Diaz-Cano, S. J., Hernández-Losa, J., & Castellví, J. (2020). Clinical implications of intratumor heterogeneity: challenges and opportunities. Journal of molecular medicine (Berlin, Germany), 98(2), 161–177. 10.1007/s00109-020-01874-2

Richardo, T., Prattapong, P., Ngernsombat, C., Wisetyaningsih, N., Iizasa, H., Yoshiyama, H., & Janvilisri, T. (2020). Epstein-Barr Virus Mediated Signaling in Nasopharyngeal Carcinoma Carcinogenesis. Cancers, 12(9), 2441. 10.3390/cancers12092441

Saliba, A. E., Westermann, A. J., Gorski, S. A., & Vogel, J. (2014). Single-cell RNA-seq: advances and future challenges. Nucleic acids research, 42(14), 8845–8860. 10.1093/nar/gku555

Saraf S, Suresh P, Das RK. Unravelling the role of EMT in OSCC: a quick peek into HPV-mediated pathogenesis. Oral Oncol Rep. 2023;5:100016.

Sberna, G., Maggi, F., & Amendola, A. (2023). Virus-Encoded Circular RNAs: Role and Significance in Viral Infections. International Journal of Molecular Sciences, 24(22), 16547. 10.3390/ijms242216547

Sertznig, H., Hillebrand, F., Erkelenz, S., Schaal, H., & Widera, M. (2018). Behind the scenes of HIV-1 replication: Alternative splicing as the dependency factor on the quiet. Virology, 516, 176–188. 10.1016/j.virol.2018.01.011

Song, H., Lim, Y., Im, H. et al. Interpretation of EBV infection in pan-cancer genome considering viral life cycle: LiEB (Life cycle of Epstein-Barr virus). Sci Rep 9, 3465 (2019). 10.1038/s41598-019-39706-0

SoRelle, E. D., Dai, J., Bonglack, E. N., Heckenberg, E. M., Zhou, J. Y., Giamberardino, S. N., Bailey, J. A., Gregory, S. G., Chan, C., & Luftig, M. A. (2021). Single-cell RNA-seq reveals transcriptomic heterogeneity mediated by host-pathogen dynamics in lymphoblastoid cell lines. eLife, 10, e62586. 10.7554/eLife.62586

SoRelle, E. D., Dai, J., Reinoso-Vizcaino, N. M., Barry, A. P., Chan, C., & Luftig, M. A. (2022). Time-resolved transcriptomes reveal diverse B cell fate trajectories in the early response to Epstein-Barr virus infection. Cell reports, 40(9), 111286. 10.1016/j.celrep.2022.111286

Swaminath S, Russell AB (2024) The use of single-cell RNA-seq to study heterogeneity at varying levels of virus–host interactions. PLOS Pathogens 20(1): e1011898. 10.1371/journal.ppat.1011898

Tagawa, T., Oh, D., Santos, J., Dremel, S., Mahesh, G., Uldrick, T. S., Yarchoan, R., Kopardé, V. N., & Ziegelbauer, J. M. (2021). Characterizing Expression and Regulation of Gamma-Herpesviral Circular RNAs. Frontiers in microbiology, 12, 670542. 10.3389/fmicb.2021.670542

Vyas, A., Harbison, R. A., Faden, D. L., Kubik, M., Palmer, D., Zhang, Q., Osmanbeyoglu, H. U., Kiselyov, K., Méndez, E., & Duvvuri, U. (2021). Recurrent Human Papillomavirus-Related Head and Neck Cancer Undergoes Metabolic Reprogramming and Is Driven by Oxidative Phosphorylation. Clinical cancer research : an official journal of the American Association for Cancer Research, 27(22), 6250–6264. 10.1158/1078-0432.CCR-20-4789.

Wu, R., Guo, W., Qiu, X., Wang, S., Sui, C., Lian, Q., Wu, J., Shan, Y., Yang, Z., Yang, S., Wu, T., Wang, K., Zhu, Y., Wang, S., Liu, C., Zhang, Y., Zheng, B., Li, Z., Zhang, Y., Shen, S., … Chen, L. (2021). Comprehensive analysis of spatial architecture in primary liver cancer. Science advances, 7(51), eabg3750. 10.1126/sciadv.abg3750

Wolpert D. Stacked generalization. Neural Networks. 1992;5:241–59.

Young, L. S., & Dawson, C. W. (2014). Epstein-Barr virus and nasopharyngeal carcinoma. Chinese journal of cancer, 33(12), 581–590. 10.5732/cjc.014.10197

Yu, X., Gong, Q., Yu, D., Chen, Y., Jing, Y., Zoulim, F., & Zhang, X. (2024). Spatial transcriptomics reveals a low extent of transcriptionally active hepatitis B virus integration in patients with HBsAg loss. Gut, 73(5), 797–809. 10.1136/gutjnl-2023-330577

zur Hausen, H., & de Villiers, E. M. (2014). Cancer "causation" by infections--individual contributions and synergistic networks. Seminars in oncology, 41(6), 860–875. 10.1053/j.seminoncol.2014.10.003

