## Supplementary Figures for "Viral Infection Detector: Ensemble Learning for Predicting Viral Infection in Single-cell Transcriptomics of Virus-Induced Cancers"

**Supplementary Figure 1. Cross validation scores for VID with XGBoost as meta-model on EBV cell line dataset**

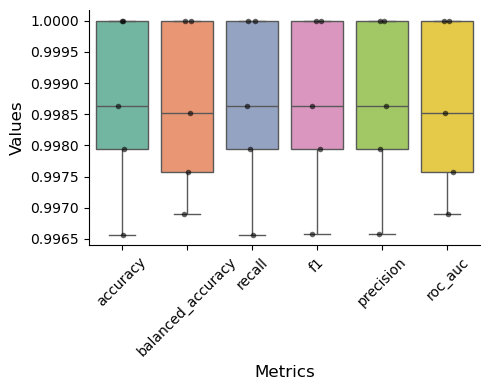

**Supplementary Figure 2. Cross validation scores for VID with XGBoost as meta-model on EBV NPC EPI dataset**

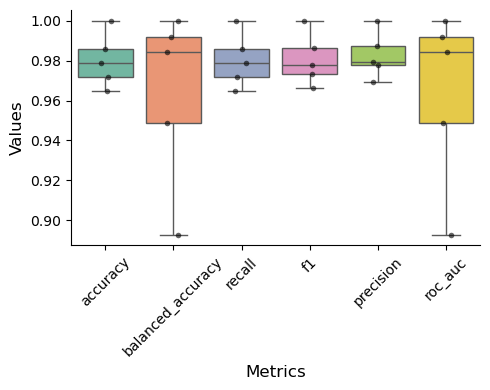

**Supplementary Figure 3. Cross validation scores for VID with XGBoost as meta-model on EBV NPC B cells dataset**

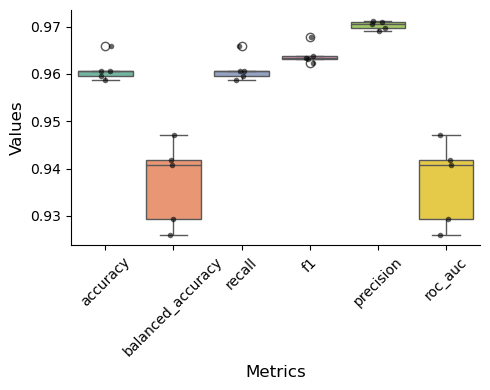

**Supplementary Figure 4. Cross validation scores for VID with XGBoost as meta-model on HPV OPC dataset**

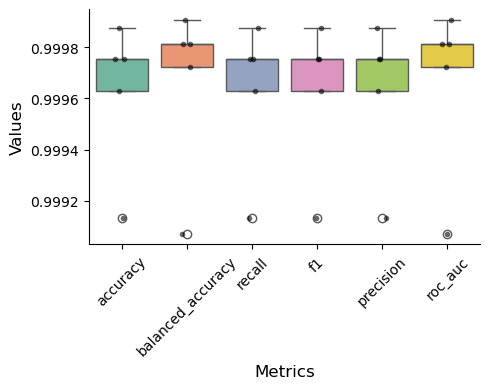

**Supplementary Figure 5. Cross validation scores for VID with MLP as meta-model on EBV cell line dataset**

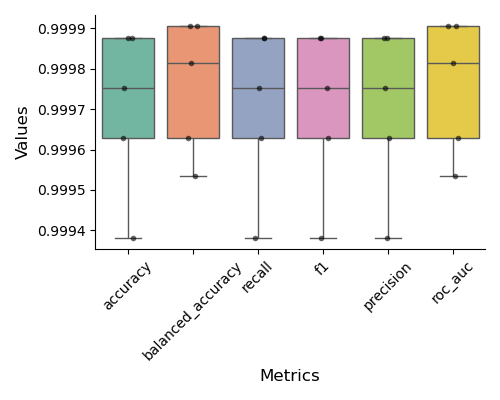

**Supplementary Figure 6.** **Cross validation scores for VID with MLP as meta-model on EBV NPC EPI dataset**

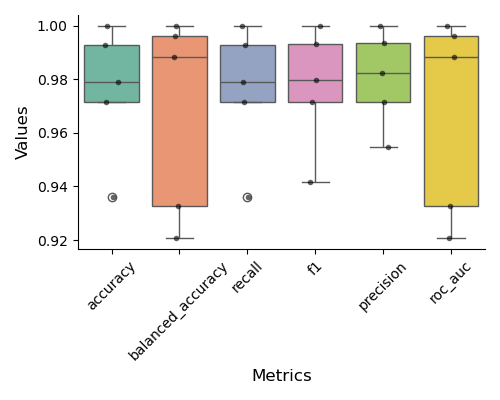

**Supplementary Figure 7. Cross validation scores for VID with MLP as meta-model on EBV NPC B cell dataset**

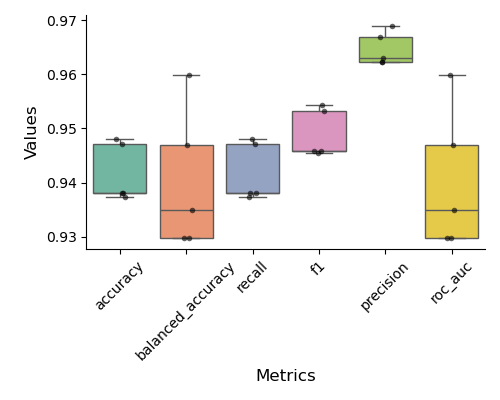

**Supplementary Figure 8. Cross validation scores for VID with MLP as meta-model on HPV OPC dataset**

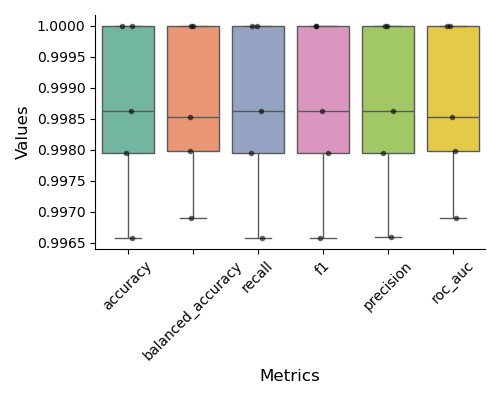

**Supplementary Figure 9. Feature importance of XGBoost for EBV cell line dataset**

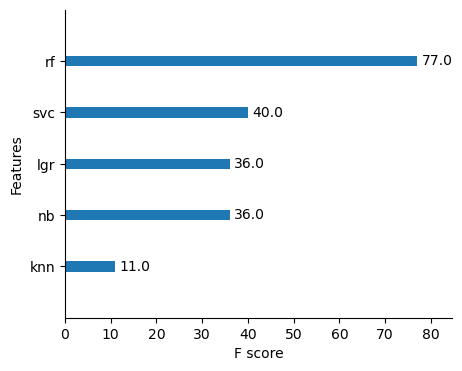
 **Supplementary Figure 10. Feature importance of XGBoost for EBV NPC EPI dataset**

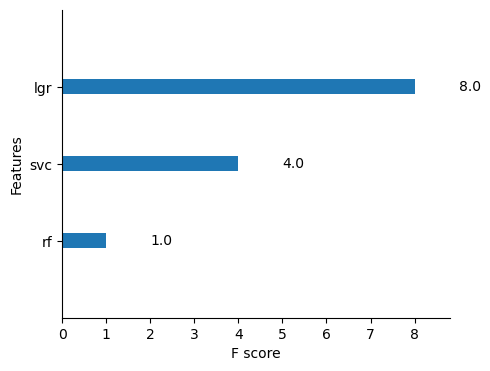

**Supplementary Figure 11. Feature importance of XGBoost for EBV NPC B cell dataset**

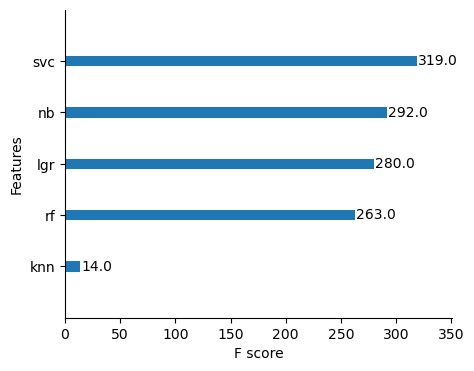

**Supplementary Figure 12. Feature importance of XGBoost for HPV OPC dataset**

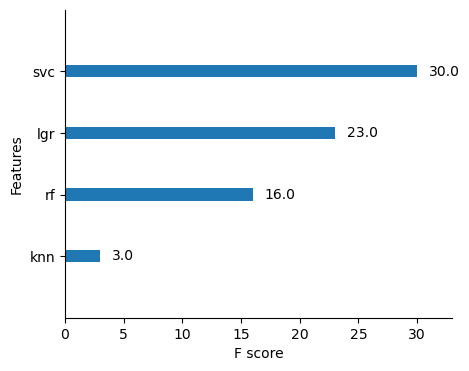

**Supplementary Table 1. Testing scores for EBV cell line dataset**

|  | **Accuracy** | **Balanced Accuracy** | **Precision** | **Recall** | **Specificity** | **F1-Score** | **AUC** |
| --- | --- | --- | --- | --- | --- | --- | --- |
| **RF** | 0.9979 | 0.9983 | 0.9979 | 0.9979 | 0.9995 | 0.9979 | 1.0000 |
| **SVC** | 0.9991 | 0.9993 | 0.9991 | 0.9991 | 1.0000 | 0.9991 | 1.0000 |
| **KNN** | 0.9977 | 0.9979 | 0.9977 | 0.9977 | 0.9986 | 0.9977 | 0.9994 |
| **GNB** | 0.9932 | 0.9937 | 0.9933 | 0.9932 | 0.9950 | 0.9932 | 0.9975 |
| **LGR** | 0.9994 | 0.9995 | 0.9994 | 0.9994 | 0.9998 | 0.9994 | 1.0000 |
| **VID(XGB)** | 0.9995 | 0.9996 | 0.9995 | 0.9995 | 0.9998 | 0.9995 | 1.0000 |
| **VID(MLP)** | 0.9995 | 0.9996 | 0.9995 | 0.9995 | 0.9997 | 0.9995 | 1.0000 |

**Supplementary Table 2. Testing scores for EBV NPC EPI dataset**

|  | **Accuracy** | **Balanced Accuracy** | **Precision** | **Recall** | **Specificity** | **F1-Score** | **AUC** |
| --- | --- | --- | --- | --- | --- | --- | --- |
| **RF** | 0.9406 | 0.7325 | 0.9404 | 0.9406 | 0.4688 | 0.9315 | 0.9814 |
| **SVC** | 0.9637 | 0.8557 | 0.9626 | 0.9637 | 0.7188 | 0.9617 | 0.9869 |
| **KNN** | 0.9538 | 0.8088 | 0.9522 | 0.9538 | 0.6250 | 0.9499 | 0.9134 |
| **GNB** | 0.9472 | 0.9291 | 0.9570 | 0.9472 | 0.9063 | 0.9503 | 0.9585 |
| **LGR** | 0.9769 | 0.9457 | 0.9773 | 0.9769 | 0.9063 | 0.9771 | 0.9874 |
| **VID(XGB)** | 0.9538 | 0.9328 | 0.9608 | 0.9538 | 0.9063 | 0.9560 | 0.9732 |
| **VID(MLP)** | 0.9670 | 0.9402 | 0.9694 | 0.9670 | 0.9063 | 0.9678 | 0.9888 |

**Supplementary Table 3. Testing scores for EBV NPC B cell dataset**

|  | **Accuracy** | **Balanced Accuracy** | **Precision** | **Recall** | **Specificity** | **F1-Score** | **AUC** |
| --- | --- | --- | --- | --- | --- | --- | --- |
| **RF** | 0.9720 | 0.7948 | 0.9725 | 0.9720 | 0.9996 | 0.9687 | 0.9807 |
| **SVC** | 0.9707 | 0.9180 | 0.9732 | 0.9707 | 0.9789 | 0.9717 | 0.9843 |
| **KNN** | 0.9632 | 0.7526 | 0.9613 | 0.9632 | 0.9960 | 0.9584 | 0.8947 |
| **GNB** | 0.9265 | 0.8309 | 0.9448 | 0.9265 | 0.9413 | 0.9335 | 0.9442 |
| **LGR** | 0.9239 | 0.9045 | 0.9556 | 0.9239 | 0.9270 | 0.9344 | 0.9638 |
| **VID(XGB)** | 0.9649 | 0.9408 | 0.9723 | 0.9649 | 0.9686 | 0.9673 | 0.9813 |
| **VID(MLP)** | 0.9398 | 0.9332 | 0.9631 | 0.9398 | 0.9409 | 0.9472 | 0.9840 |

**Supplementary Table 4. Testing scores for HPV OPC dataset**

|  | **Accuracy** | **Balanced Accuracy** | **Precision** | **Recall** | **Specificity** | **F1-Score** | **AUC** |
| --- | --- | --- | --- | --- | --- | --- | --- |
| **RF** | 0.9949 | 0.9954 | 0.9949 | 0.9949 | 0.9974 | 0.9949 | 0.9996 |
| **SVC** | 0.9974 | 0.9980 | 0.9975 | 0.9974 | 1.0000 | 0.9974 | 1.0000 |
| **KNN** | 0.9901 | 0.9872 | 0.9902 | 0.9901 | 0.9763 | 0.9901 | 0.9983 |
| **GNB** | 0.9837 | 0.9862 | 0.9842 | 0.9837 | 0.9956 | 0.9837 | 0.9957 |
| **LGR** | 0.9974 | 0.9978 | 0.9975 | 0.9974 | 0.9991 | 0.9974 | 1.0000 |
| **VID(XGB)** | 0.9987 | 0.9990 | 0.9987 | 0.9987 | 1.0000 | 0.9987 | 1.0000 |
| **VID(MLP)** | 0.9978 | 0.9982 | 0.9978 | 0.9978 | 1.0000 | 0.9978 | 1.0000 |
